## Supplementary file with optimization model details for "Elastic energy savings and active energy cost in a simple model of running"

### S1 Appendix for "Elastic energy savings and active energy cost in a simple model of running". Dynamic optimization model details.

**Ryan T. Schroeder[1] and Arthur D. Kuo[1,2]**

**[1]Faculty of Kinesiology, University of Calgary, Alberta, Canada**

**[2]Biomedical Engineering Program, University of Calgary, Alberta, Canada**

We provide additional details regarding the derivation of model equations and implementation of the dynamic optimization problem. The dynamic optimization was used to determine optimal actuation strategies for running over multiple speeds on flat and sloped ground. The simple model constitutes a point-mass body representing the center of mass (CoM) of the organism and a massless leg consisting of a compression spring connected in series with a length actuator that can actively extend or resist compression (Fig. 2c of the main text). The model's leg neglects more complex morphology associated with an animal leg. However, the spring and length actuator conceptually represent gross compliance associated with various passive (tendon) and active (muscles) tissues utilized during running. Two modes of passive energy dissipation were included in the model: collision loss associated with foot-ground contact and hysteresis of the tendon.

The collision was formulated as a discontinuous reduction in the CoM velocity vector magnitude at touchdown. The collision fraction CF was purely absorptive and did not affect the CoM velocity direction. The resulting fraction of lost kinetic energy was given by the change in the CoM velocity magnitude from before ($v^-$) to after ($v^+$) the collision: $v^+ = v^-(1 - \mathrm{CF})$.

$$\text{collision loss} = \frac{\frac{1}{2}M(v^+)^2 - \frac{1}{2}M(v^-)^2}{\frac{1}{2}M(v^-)^2} = (\mathrm{CF}^2 - 2\mathrm{CF}) \qquad \text{(s1)}$$

Where $M$ is body mass and nominal CF is 0.03, resulting in about 6% loss. Tendon hysteresis was modeled as a viscous damper operating in parallel with the model's spring, active only during compression. The damping coefficient $c$ was calculated with damping ratio $\zeta$, where $c = 2\zeta\sqrt{kM}$, and $k$ is the spring stiffness. The model was optimized for a damping value of $\zeta = 0.1$. The energy loss due to hysteresis was given by

$$\text{hysteresis loss} = 1 - \exp\left(-4\zeta \frac{\cos^{-1}\left(\sqrt{\frac{1+\zeta}{2}}\right)}{\sqrt{1-\zeta^2}}\right) \qquad \text{(s2)}$$

which yielded about 26% loss. The spring stiffness was explored over a range of values: $4.93 \leq k \leq 122\ \mathrm{kN \cdot m^{-1}}$. These numbers were chosen to encompass a comprehensive spectrum of gait ranging from “grounded running” solutions, where no flight phase occurs due to an extremely compliant spring, to “impulsive running,” where a brief stance period is followed by a long flight phase. The limiting case of infinitesimal stance due to an infinitely stiff spring was also considered, conceptually (i.e. not through optimization). In this case, ballistics equations were used to simulate the body’s parabolic trajectory through the air and touchdown/takeoff velocities were assumed to occur with equal magnitude and opposite vertical direction.

Step length $s$ was implemented as a constraint on the horizontal distance travelled by the CoM over a step. Step frequency $f$ was applied as a constraint on the time duration of the step ($T = 1/f$). Together, these constraints enforced the desired running speed in each simulation. Gait was optimized over a range of speeds $2 - 4\ \mathrm{m \cdot s^{-1}}$, while step frequency was determined by empirical trends of treadmill running ($f = 0.26v + 2.17$; (1)) and step length was calculated with $s = v/f$.

The model assumes no slipping at the foot-ground contact point $x_f$, included as a decision variable. A single-phase optimization was utilized to simulate ground contact (stance phase) of a single running step and boundary conditions were determined via kinematics derived from ballistics of a point mass during no ground contact (flight phase). The optimization output of a single step was assumed to repeat in sequence with the contralateral leg (i.e. symmetrical gait) and periodic gait was enforced.

The state vector for the model is given.

$$\mathbf{q} = [x_b \quad \dot{x}_b \quad y_b \quad \dot{y}_b \quad L_m \quad \dot{L}_m \quad \ddot{L}_m]^T \qquad \text{(s3)}$$

Where $x_b$ and $y_b$ are the tangential and normal displacement of the CoM along the substrate and $\dot{x}_b$ and $\dot{y}_b$ are their respective time derivatives. $L_m$, $\dot{L}_m$ and $\ddot{L}_m$ are the length, extension velocity and acceleration of the actuator, respectively. The state derivative is given.

$$\dot{\mathbf{q}} = [\dot{x}_b \quad \ddot{x}_b \quad \dot{y}_b \quad \ddot{y}_b \quad \dot{L}_m \quad \ddot{L}_m \quad \dddot{L}_m]^T \qquad \text{(s4)}$$

Where,

$$\ddot{x}_b = \frac{x_b - x_f}{ML_l}(F_s + F_d) - g \sin\alpha \tag{s5}$$

$$\ddot{y}_b = \frac{y_b}{ML_l}(F_s + F_d) - g \cos\alpha \tag{s6}$$

In Eqs. (s5) & (s6), $F_S$ and $F_d$ are spring and damping forces of the tendon, $L_l$ is the total length of the leg as a function of time, $g$ is the gravitational acceleration constant ($9.81\ \mathrm{m \cdot s^{-2}}$) and $\alpha$ is the ground slope angle relative to horizontal (positive indicating upward slope).

$$F_s = k\left(L_{t,o} - L_t\right) \tag{s7}$$

$$F_d = \begin{cases} c\dot{L}_t, & \dot{L}_t \geq 0 \\ 0, & \dot{L}_t < 0 \end{cases} \tag{s8}$$

$$L_l = \sqrt{\left(x_b - x_f\right)^2 + y_b^2} \tag{s9}$$

Where $L_t$ is the instantaneous length of the spring and $L_{t,o}$ is the equilibrium length of the spring associated with zero spring force. Leg length was constrained so as not to exceed a maximum allowable value, $L$. Given the serial configuration of the actuator-spring unit, $L_l = L_t + L_m$ and $F_m = F_s + F_d$, where $F_m$ is the muscle actuator force. The vector array, $\mathbf{u}$, contains the control variables implemented in the optimization problem.

$$\mathbf{u} = [\dddot{L}_m \quad P_m^+ \quad P_m^-]^T \tag{s10}$$

Where $\dddot{L}_m$ is jerk of the leg's extension and $P_m^+$ and $P_m^-$ are slack variables used to capture positive and negative components of the muscle actuator's power, given the path constraint, $P_m = P_m^+ + P_m^-$ where $P_m = F_m\dot{L}_m$. When damping was included in the model, $\mathbf{u}$ was expanded to include slack variables for the positive and negative components of damper's velocity so as to implement damping forces during compression only.

$$\mathbf{u} = [\dddot{L}_m \quad P_m^+ \quad P_m^- \quad \dot{L}_t^+ \quad \dot{L}_t^-]^T \tag{s11}$$

Complementarity constraints were enforced by including $\epsilon_o P_m^+ P_m^-$ and $\epsilon_o \dot{L}_t^+ \dot{L}_t^-$ in the objective function so as to avoid inappropriate simultaneous positive and negative actuator power or damper velocity in the model. In all optimizations, these terms were driven toward zero and did not contribute to the cost of the model.

The objective function (Eq. 1, main text) included actuator work per step ($W_m$), converted to energy by scaling with positive and negative efficiencies of muscle (Eq. 2, main text). The force-rate cost was formulated as integration of the force rate squared over a single step (Eq. 3, main text). Models were parametrically evaluated with a range of force-rate cost coefficients $\epsilon$, including zero meaning a work-only cost.

The model was implemented with and without individual components. For example, it was tested without elastic energy return (i.e. no spring, or infinite spring stiffness), and without dissipation as the Actuator-only model. The Actuated spring-mass model includes all of the components, but with parameters such as collision fraction and hysteresis varied in sensitivity analyses.

All optimizations were conducted with the MATLAB software GPOPS-II (2) and the resulting nonlinear problem was solved using SNOPT (3). First, fifteen random initial guesses were used to sample multiple local optima over the solution space, and then the lowest-cost solution was repeatedly perturbed (thirty times) with 8% random noise (scaled to all states and controls) in order to fine tune the presumed global optima (procedure from (4)). All variables and equations were non-dimensionalized with parameter combinations ($L$, $M$ and $g$) during optimization and outputs were subsequently re-dimensionalized as indicated in figures.
